# The Nutrient and Energy Pathway Requirements for Surface Motility of Nonpathogenic and Uropathogenic *Escherichia coli*

**DOI:** 10.1101/2020.08.12.249037

**Authors:** Sushmita Sudarshan, Jacob Hogins, Sankalya Ambagaspitiye, Philippe Zimmern, Larry Reitzer

**Author notes:** Address correspondence to Larry Reitzer.

## Abstract

Uropathogenic *E. coli* (UPEC) is the causative pathogen for most uncomplicated urinary tract infections. Flagellar-mediated motility is essential for virulence and colonization for ascending urinary tract infections. The appendage requirement for surface motility depends on the strain: nonpathogenic *E. coli* (NPEC) lab strains use pili, NPEC hypermotile derivatives use flagella, and UPEC strains use flagella. *E. coli* flagella-dependent surface motility had been previously shown to require glucose and amino acids. We examined the nutritional and pathway requirements of the NPEC strain W3110 for pili-dependent surface motility, which have not been previously examined. We then compared these requirements to those for two strains with flagella-dependent surface motility: a variant of W3110, W3110-J1, in which the synthesis of the activator of flagella synthesis has been upregulated and the UPEC strain UTI89. The glucose requirement for W3110 was higher than that for either W3110-J1 or UTI89. The pathways required for motility were also different. W3110, but not UTI89, required the Embden-Meyerhof-Parnas pathway via PfkA; conversely, UTI89, but not W3110, required the Entner-Doudoroff pathway, acetogenesis, and the TCA cycle. Glucose did not control flagella synthesis for W3110-J1 and UTI89. The differing requirements for surface motility are likely to reflect major metabolic differences between strains. The metabolic requirements for UTI89 motility suggest a specific adaptation to the urinary tract environment.

**IMPORTANCE:** Urinary tract infections affect over 50% of women and *E. coli* is the most common uropathogen. Virulence requires both pili and flagella, and both appendages can contribute to surface motility. Previous studies of *E. coli* surface motility did not consider the appendage requirement and the ability to switch the surface appendage. The nutrient and pathway requirements for surface motility of a non-pathogenic *E. coli* strain with either pili- or flagella-dependent surface motility and the uropathogen UTI89 were examined. Pili-dependent surface motility required glycolysis, while flagella-dependent motility required the TCA cycle and oxidative phosphorylation and was less dependent on glycolysis. The distinctive nutrient and pathway requirements for UTI89 motility probably result from metabolic adaptations to the urinary tract.

## INTRODUCTION

Swarming is flagella-dependent bacterial surface motility (1-3). Swarming cells express virulence genes and show enhanced resistance to both engulfment and antibiotics (4-7). The genetic requirements for *E. coli* swarming were examined for mutants of the Keio collection, which contains deletions in most non-essential genes (8). Swarming motility required flagella, but unexpectedly also required pili. The authors suggested that some genes for pili synthesis were required for flagella synthesis which is not consistent with subsequent evidence that pili and flagella synthesis are mutually exclusive (9-11). Our recent results show that surface motility of common *E. coli* lab strains, including the parental strain of the Keio mutants, requires pili, but fast variants rapidly appear due to a mutation that increases expression of the master regulator for flagella synthesis (11). In other words, results using Keio collection mutants involved strains that either expressed pili or flagella or both if a flagella-synthesizing variant was generated during the motility assay. The requirements for motility are likely to depend on the appendage and must be reexamined in strains for which the motility appendage is unambiguously known.

Urinary tract infections (UTIs) are one of the most common bacterial infections, affecting approximately 150 million people worldwide each year (12). UTIs have produced an increasing burden on the healthcare system because of recurrence and antibiotic resistance (13). Women are more prone to UTIs than men with over 50% of women experiencing at least one infection in their lifetime (13). The most common uropathogen is *E. coli* (14, 15), which is responsible for about 80-90% of community acquired UTIs and 40-50% of nosocomial acquired UTIs (13). A recent study found that *E. coli* was present in the urine of 65.5% of 4453 women with UTIs (16).

*E. coli* generally resides in the intestinal tract, but uropathogenic *E. coli* (UPEC), a pathotype of extra intestinal pathogenic *E. coli*, can migrate, adapt and colonize the urinary tract and cause a urinary tract infection (UTI) (17). UPEC can infect the urinary tract, kidneys, and bloodstream causing cystitis, pyelonephritis, and sepsis, respectively. Nutrient availability differs between the intestinal and urinary tracts (18, 19). The metabolic pathways required for growth in each environment differ and UPEC metabolism is adapted to the urinary tract. For example, growth of *E. coli* in the intestinal tract requires multiple carbohydrates, while growth in the urinary tract is proposed to require amino acids and peptides (18, 20, 21).

A UTI also requires flagella-mediated movement (22). In a murine model, the UPEC strain CFT073 required flagellin, the *fliC* product, to ascend to the upper urinary tract (22). 95% of UTIs may be ascending infections meaning that the infection begins by colonization of the periurethral area, followed by movement up the urethra into the bladder, and possibly into the ureters and kidneys (23). The implicit assumption of such studies is that movement only requires flagella. However, the presentation of human UTI varies from localized trigonitis to unlocalized pancystitis (24). The movement required for establishment of a localized UTI may differ from that required for progression to pancystitis and the latter could conceivably involve pili, even though pili are primarily associated with adhesion.

Our goals were to examine the nutrient and pathway requirements for surface motility of a non-pathogenic *E. coli* lab strain with either pili- or flagella-dependent surface motility, and to compare these requirements to those of the UPEC strain UTI89. We then discuss our results in relation to nutrient and pathway requirements for CFT073 infection in a mouse model.

## RESULTS

### Requirement of glucose for surface motility

We examined the glucose requirement for surface motility for three strains: non-pathogenic W3110 which requires pili for surface movement; J1 which is a hypermotile derivative of W3110 that utilizes flagella for surface movement because of an IS5 insertion in the *flhDC* promoter region; and uropathogenic UTI89 which also moves with flagella (11). Electron microscopic images of these strains taken from surface motility plates confirmed their appendage requirement (Fig 1). W3110 moved relatively slow, did not reach the plate’s edge during the assay, and formed a ring pattern which resembles the swarm pattern of *Proteus mirabilis* (25) (Fig 2). Its movement required 0.5% glucose. J1 and UTI89 movement with 0.5% glucose covered the entire plate with no discernible pattern. J1 moved less well with 0.25% glucose, but UTI89 moved normally. J1 and UTI89 did not move with 0.125% glucose. The glucose requirement was different for each strain, and the strains with flagella-dependent surface motility had a lower glucose requirement.

**Figure 1:**
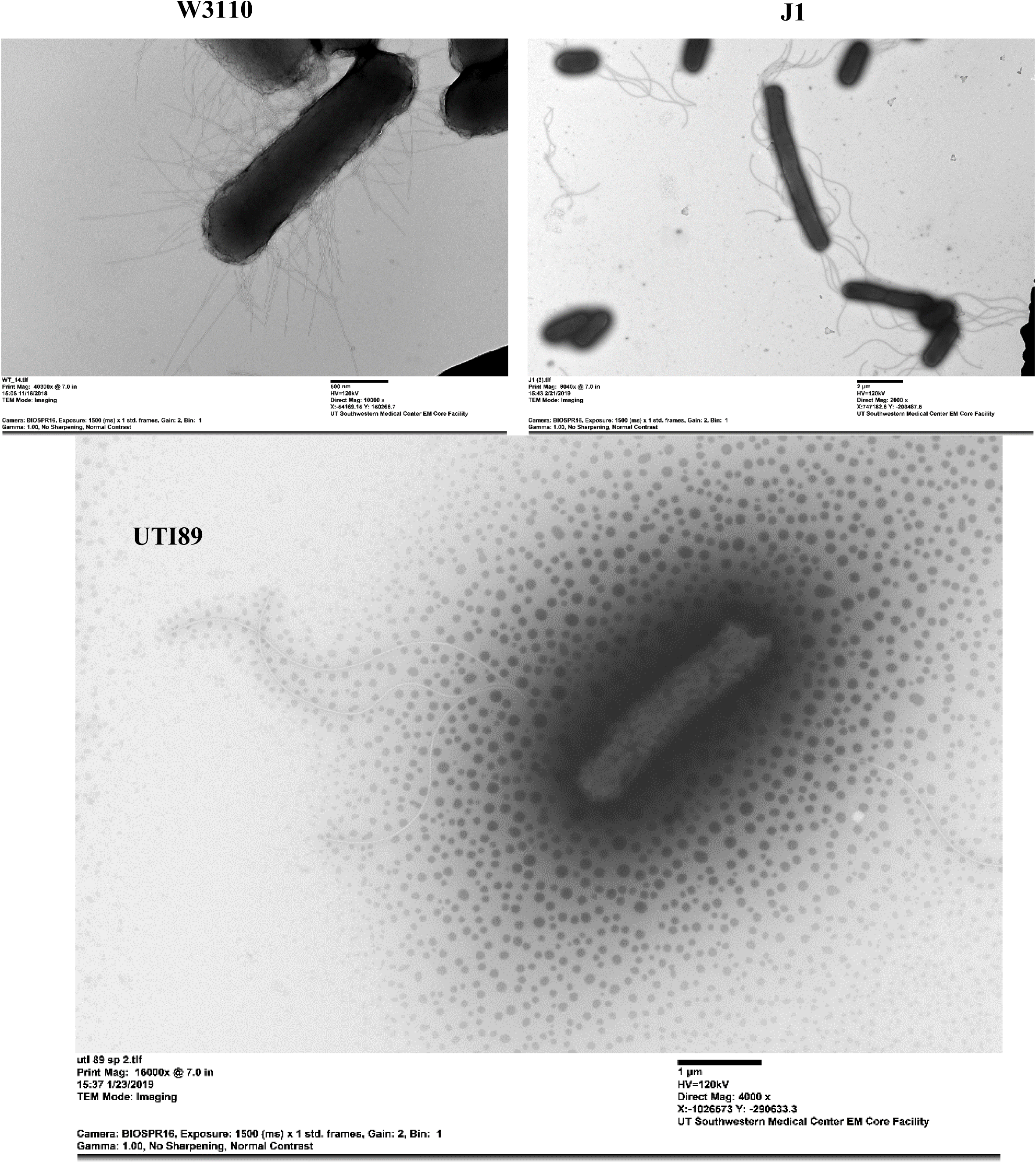
Electron microscopic images of nonpathogenic W3110 and J1, and uropathogenic UTI89. UTI89 and J1 possess flagella which are longer than the cell length, while W3110 possess pili. The dots around UTI89 are typical for EM images of this strain and probably indicate the presence of exopolysaccharide. The bars for W3110, J1 and UTI89 are 0.5, 2, and 1 µm, respectively.

**Figure 2:**
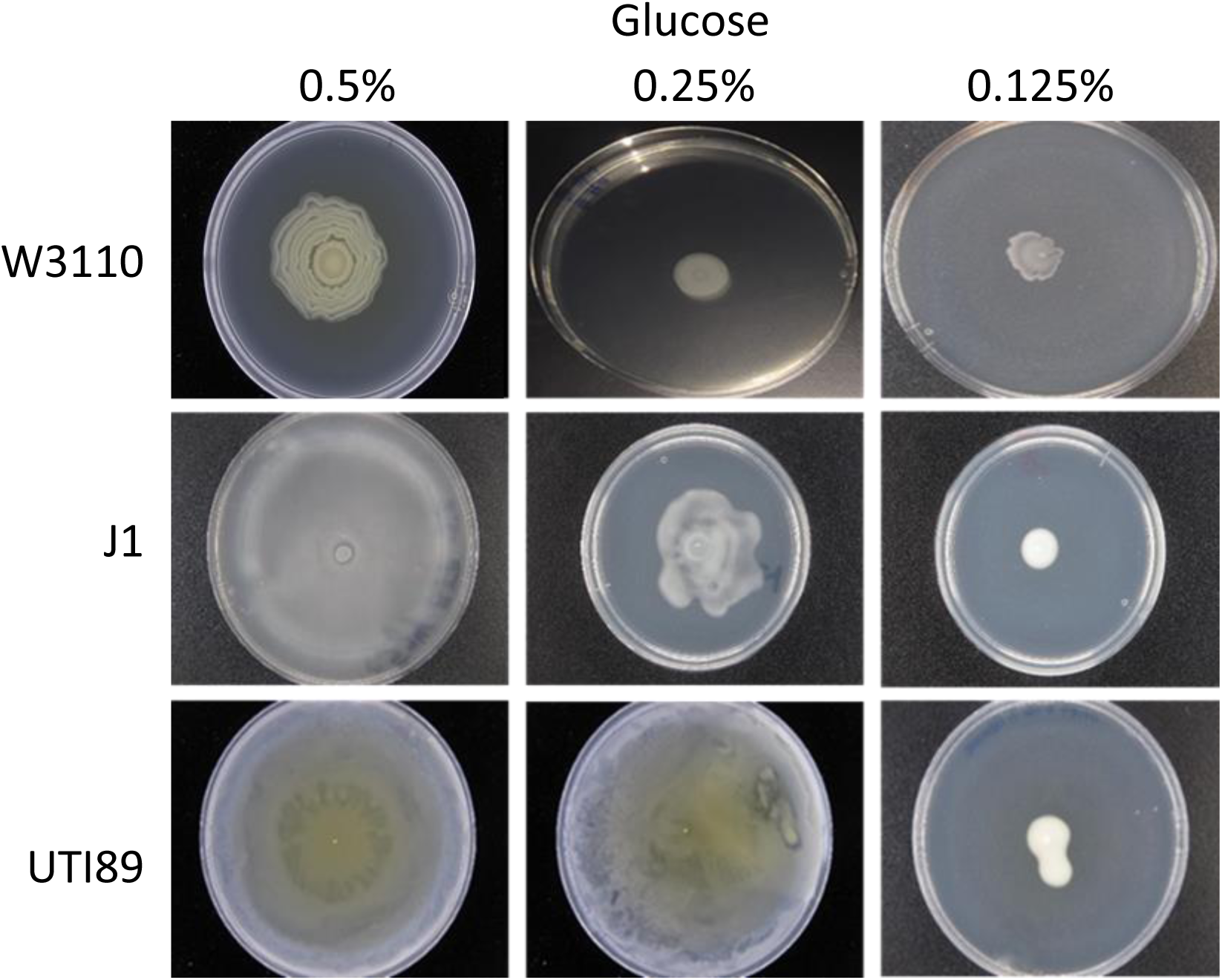
Concentration requirement of glucose for surface motility.

### Surface-motility of mutants with defects in glucose transport and pathways of carbohydrate metabolism

We examined surface motility in mutants with defects in glucose transport and the following pathways of central metabolism: the Embden-Meyerhof-Parnas (EMP) pathway, which is the standard glucose-degrading pathway for numerous organisms including *E. coli*; gluconeogenesis; the oxidative branch of the pentose cycle; the Entner-Doudoroff (ED) pathway, which is an alternate glycolytic pathway that degrades glucose to pyruvate; and acetogenic enzymes that convert pyruvate to acetate. Mutants with defects in these pathways have been examined in mouse models of UTIs (reviewed in (18)). These pathways are important for energy generation and biosynthesis of a variety of intermediates, such as NADPH and ribose-5-phosphate, triose phosphates for the glycerol backbone of phospholipids, 3-phosphoglycerate for serine, glycine, and cysteine synthesis, pyruvate for acetyl-CoA synthesis, and acetyl-CoA for either the TCA cycle or acetate formation.

The results of the mutational analysis are shown in Figs 3 and S1 and summarized in Table 1. For W3110, surface motility required a gene for the rate-limiting enzyme of the EMP pathway, *pfkA* (phosphofructokinase); three genes of the major glucose transport system — *ptsG, ptsH, ptsI*; the first gene of the oxidative pentose cycle, *zwf* (glucose-6-phosphate dehydrogenase); and a gene required for synthesis of the 3-phosphoglycerate family of amino acids, *serB* (phosphoserine phosphatase). W3110 moved less well with deletions in genes of gluconeogenesis, *pck* (phosphoenolpyruvate carboxykinase) and the ED pathway, *edd* (phosphogluconate dehydratase) and *eda* (KDPG aldolase). Loss of acetogenic genes — *pta* (phosphotransacetylase) and *ackA* (acetate kinase) — did not impair movement.

**Table 1:** Summary of surface motility of mutants with defects in various pathways. a The scoring is ++ for ≫ 70% of parental diameter; + for 30-50% of parental diameter; and — for < 25% of parental diameter. Diameter assessments are qualitative because of variations from batch to batch of motility assay plates. b The results from a mouse infection model have been reviewed (18). c These exact mutants were not tested in a mouse infection, but a deletion in the same operon had the indicated result. ND, not determined.

| Genotype | Pathway affected | W3110 <sup>a</sup> | J1 <sup>a</sup> | UTI89 <sup>a</sup> | Infection model <sup>b</sup> |
| --- | --- | --- | --- | --- | --- |
| parental | none | ++ | ++ | ++ | + |
| $\Delta ptsI$ | Glucose transport | — | ++ | + | ND |
| $\Delta ptsH$ | Glucose transport | — | ++ | + | ND |
| $\Delta ptsG$ | Glucose transport | — | ++ | ++ | ND |
| $\Delta pfkA$ | EMP glycolysis | — | — | ++ | + |
| $\Delta edd$ | ED glycolysis | + | ++ | — | + |
| $\Delta eda$ | ED glycolysis | + | ++ | + | ND |
| $\Delta zwf$ | Oxidative pentose cycle | — | ++ | + | + |
| $\Delta pta$ | Acetogenesis | ++ | ++ | — | — |
| $\Delta ackA$ | Acetogenesis | ++ | ++ | + | — |
| $\Delta pck$ | Gluconeogenesis | + | ++ | ++ | — |
| $\Delta serB$ | Serine synthesis | — | ++ | — | ND |
| $\Delta gltA$ | TCA cycle | + | + | — | ND |
| $\Delta sdhA$ | TCA cycle | + | + | — | — <sup>c</sup> |
| $\Delta sucC$ | TCA cycle | + | + | — | — <sup>c</sup> |

**Figure 3:**
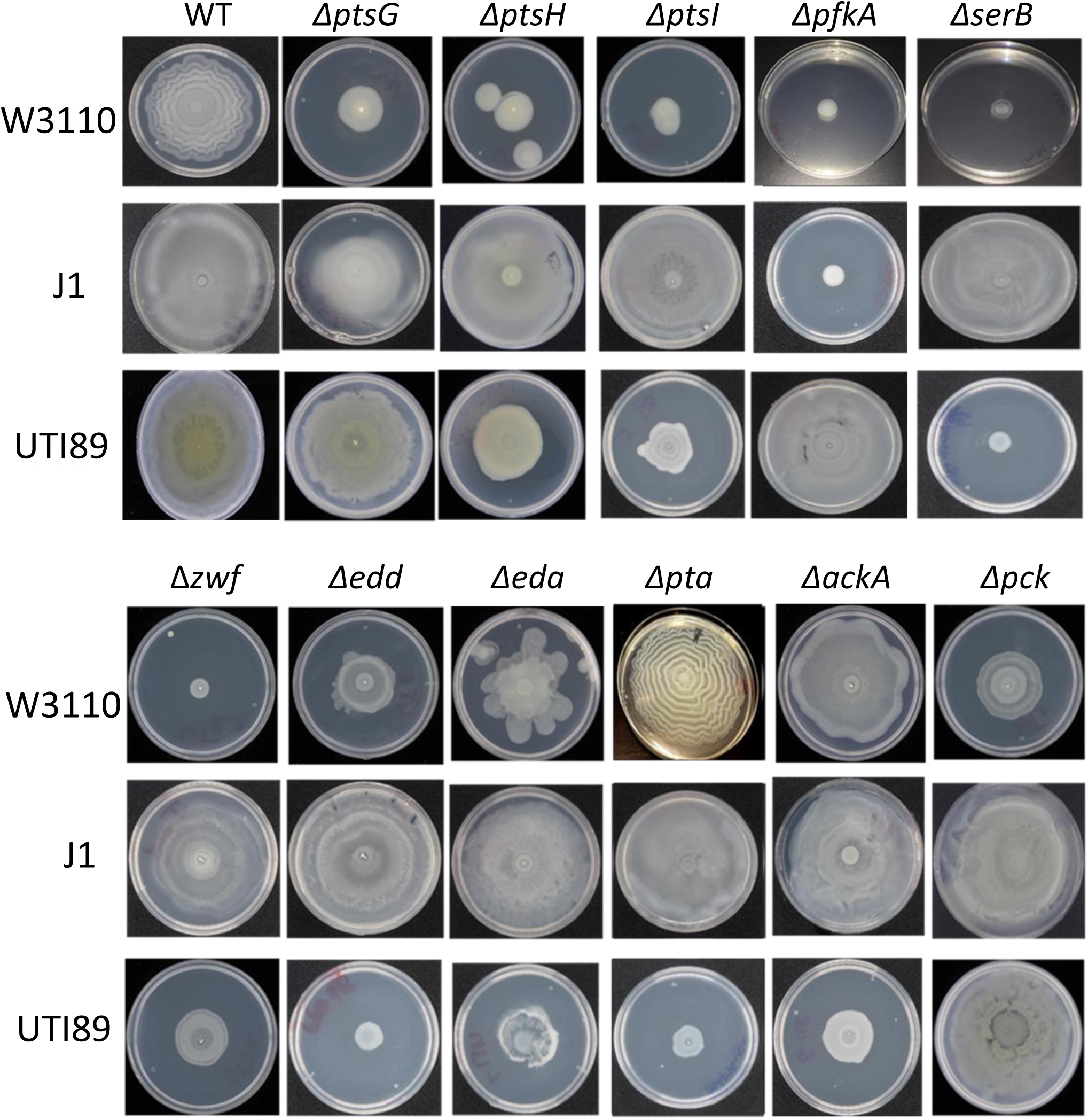
Surface motility of mutants with defects in glucose transport and pathways of carbohydrate metabolism.

J1 surface motility required PfkA, but unlike parental W3110 did not require the oxidative pentose cycle (*zwf*) or components of the major glucose transport system—*ptsG, ptsH*, and *ptsI*.

The pathway requirements for pathogenic UTI89 were different from both W3110 and J1. UTI89 movement was not affected by loss of *pfkA, ptsG*, and *pck*, and was substantially or completely impaired by loss of *ptsI, ptsH*, the ED pathway genes *eda* and *edd, zwf*, the acetogenic genes *ackA* and *pta*, and *serB*.

### Factors that affect the glucose requirement

#### Iron limitation

Iron can modulate the expression of genes of energy metabolism, so we examined the effect of supplemental 2,2-bipyridyl (BIP), an iron chelator, on surface motility (26). With BIP (low iron) UTI89 moved with 0.125% glucose (Fig 4), but not without glucose (not shown). Low iron restored movement of J1 with 0.25% glucose and allowed partial movement with 0.125% glucose. W3110 showed some movement with 0.25% glucose but no movement at 0.125% glucose (Fig 4). Surface motility plates with BIP and no glucose did not support growth in any of the strains (not shown). Even with low iron, flagella-mediated motility required a lower concentration of glucose than pili-mediated motility.

**Figure 4:**
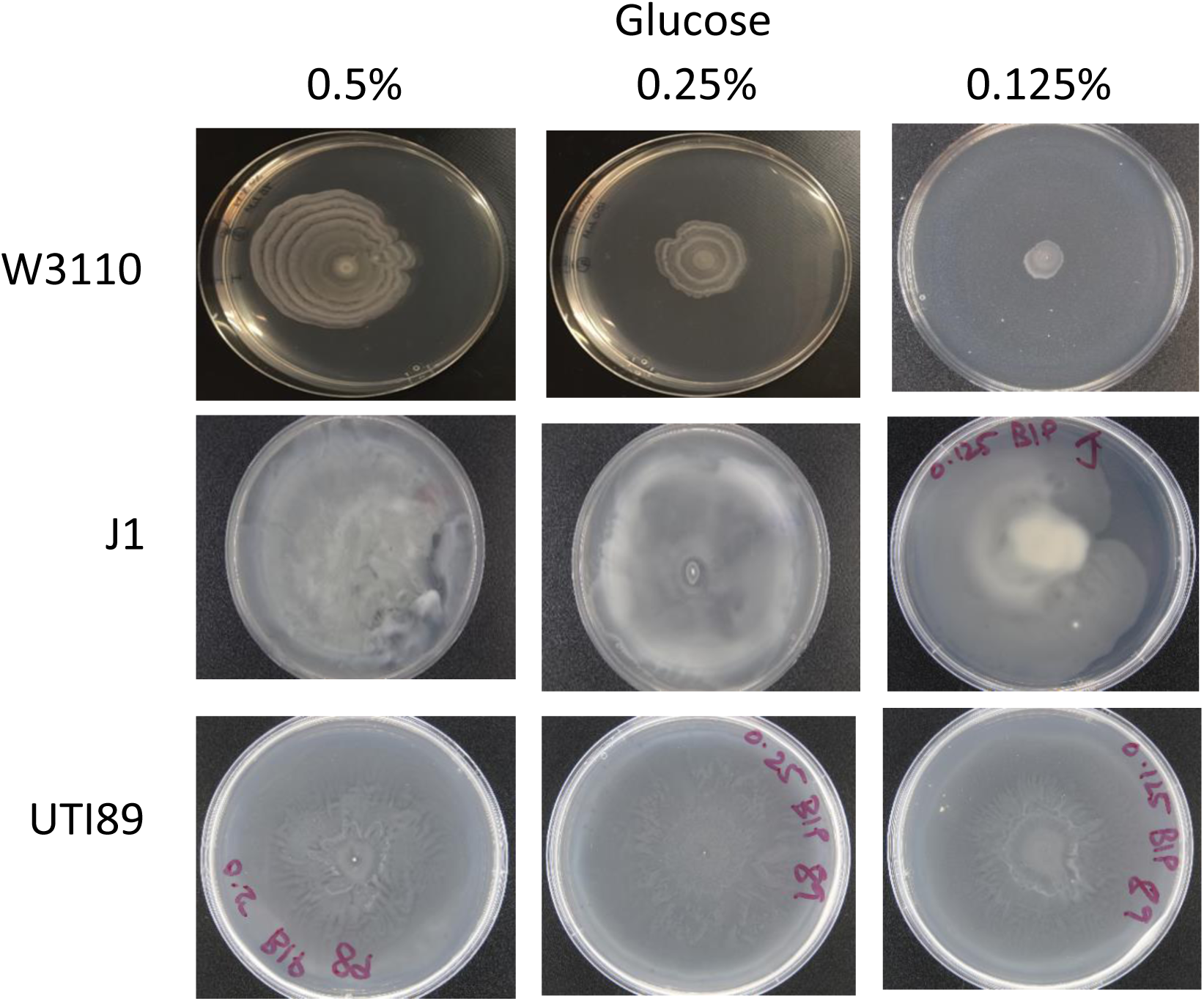
Surface motility of W3110, J1 and UTI89 with 100 µM BIP and the indicated concentrations of glucose.

#### Glycerol, cysteine, and pyruvate

We examined whether glycerol, cysteine, and pyruvate could reduce the glucose requirement for W3110 because: (a) we could not genetically test the importance of glycerol-3-P which is required for phospholipid synthesis, (b) a higher than normal level of cysteine has been shown to be a requirement for *S. enteric* surface motility (27, 28), which is consistent with the result that a W3110 Δ*serB* mutant failed to move (Fig. S1), and (c) pyruvate is a product of carbohydrate catabolism and is a major branch point of metabolism that can provide energy via the TCA cycle, and generate alanine and acetate.

Supplements were added to medium with 0.125% glucose and 100 µM BIP, which does not support surface motility of W3110 (Fig 4). A combination of 2 mM cysteine, 0.1% glycerol, and 0.1% pyruvate stimulated surface motility (Fig 5). Cysteine omission did not prevent movement but altered the motility pattern (Fig 5); glycerol omission severely impaired movement (Fig 5); and pyruvate omission had no effect on movement (Fig 5). Glycerol alone stimulated movement, but cysteine did not (Fig 5). Glycerol replaced glucose for strains J1 and UTI89 (Fig 6). In summary, glycerol lowered the glucose requirement, and could replace glucose for the two strains with flagella-mediated surface motility.

**Figure 5:**
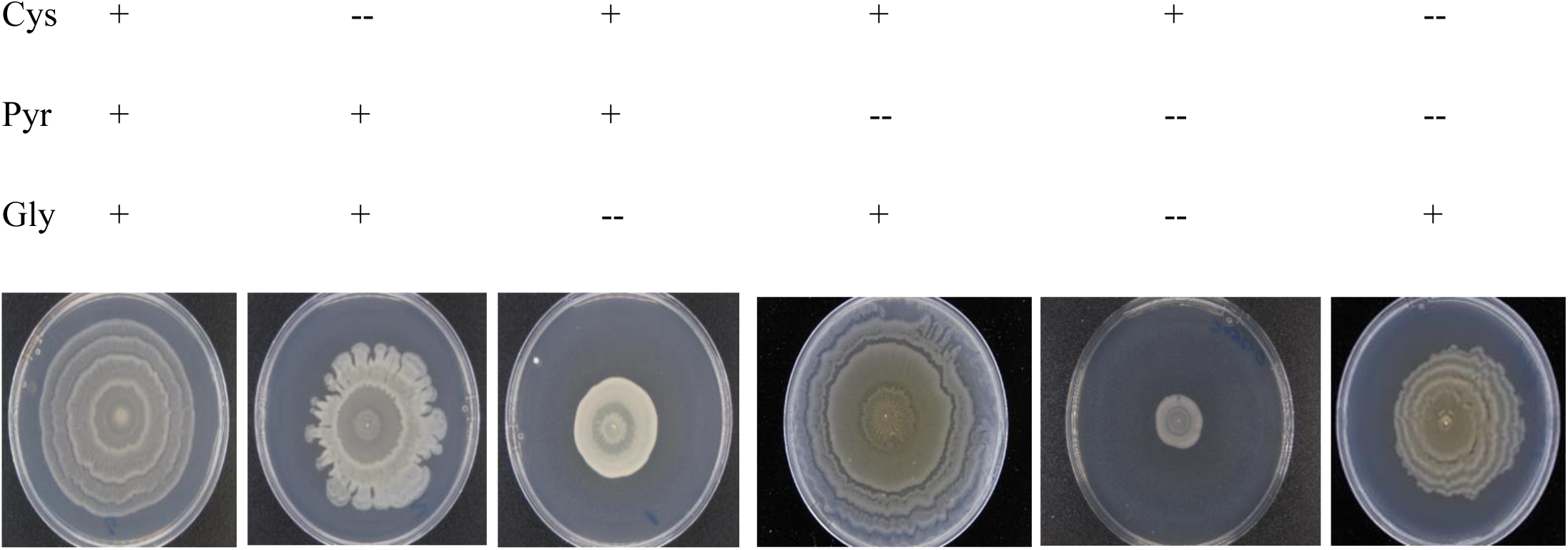
W3110 surface motility with cysteine, pyruvate and glycerol. The medium contained 0.125% glucose and 100 µM BIP. Cys, 2 mM cysteine; Pyr, 0.1% pyruvate; Gly, 0.1% glycerol.

**Figure 6:**
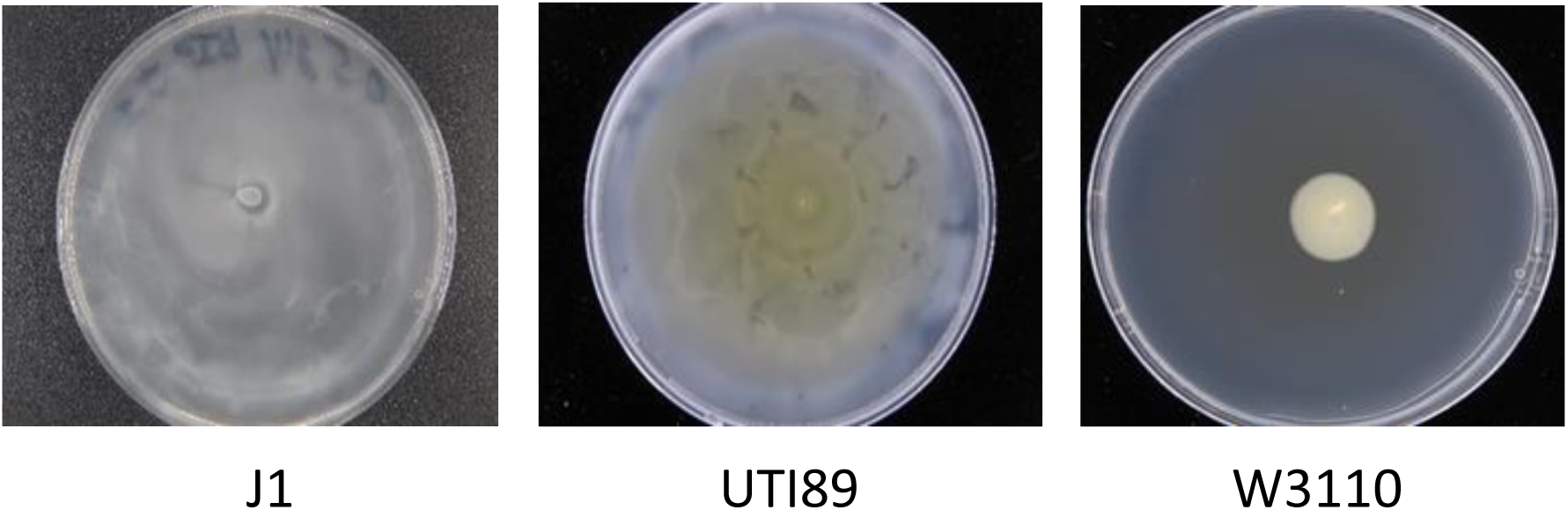
Surface motility with 0.5% glycerol instead of 0.5% glucose.

#### Supplementation with other sugars

Our results suggest that surface motility requires a carbohydrate (Fig 2). Glucose and glycerol are not abundant in urine, but urine does contain carbohydrates. Single urinary carbohydrates are under 0.4 mM, except for glucuronate (∼2 mM in urine), but the total urinary carbohydrate content is substantial (>4 mM) (9). A mixture of glucuronate (5 mM), gluconate (1 mM), glycerol (1 mM), glucose (1 mM), mannitol (1 mM), and sorbitol (1 mM) did not support the motility of strain J1 but supported motility of UTI89 weakly after 24 hours (the inner circle in Fig 7A). The appearance of a hypermotile uropathogenic variant was apparent after 24 hours, and the variant moved to the edge of the plate after 48 hours (Fig 7B). Cells from the edge of the plate were isolated and shown to move without any carbohydrate (Fig 7C). A possible explanation for enhanced motility is an insertion in the promoter region of the *flhDC* operon (11, 29). PCR analysis of this region showed no insertion upstream from the *flhDC* structural genes (not shown). These variants were not further characterized. In summary, motility without a carbohydrate is a property of a UTI89 variant, not a property of parental UTI89.

**Figure 7:**
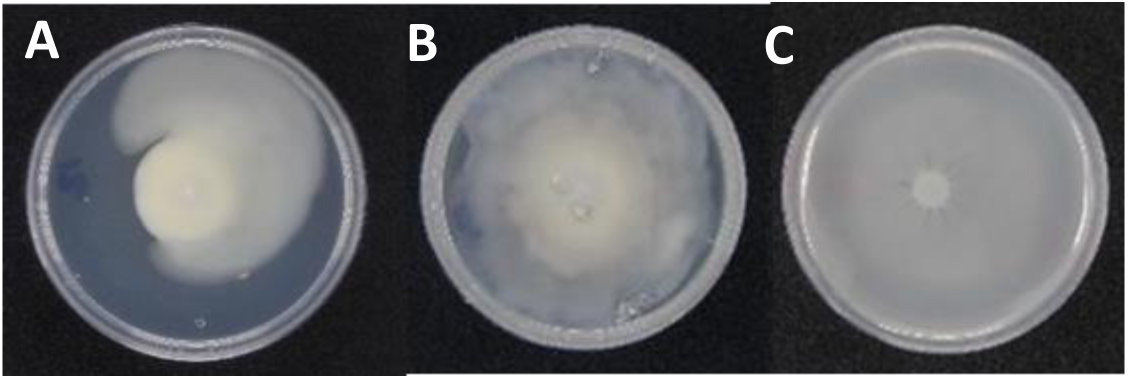
Surface motility of UTI89 with a carbohydrate mixture. The carbohydrate mixture contained 5 mM glucuronate, 1 mM gluconate, 1 mM glucose, 1 mM glycerol, 1 mM mannitol, and 1 mM sorbitol. (A) Motility for 24 hours with the carbohydrate mixture. Notice the presence of faster moving flares. (B) Motility for 48 hours with the carbohydrate mixture. (C) Cells from edge of plate B were incubated for 15 hours in the absence of any carbohydrate.

### Glucose is required for type I pili-mediated surface motility

We observed surface motility of W3110 on plates with 0.25% agar, which is used to monitor swimming motility, but only if the medium contained glucose (Fig 8). In the absence of glucose, W3110 swam into plates containing 0.25% agar, and such movement required flagella (FliC) (Fig 8). In the presence of glucose, W3110 moved on the surface, and this movement required pili (FimA) (Fig 8).

**Figure 8:**
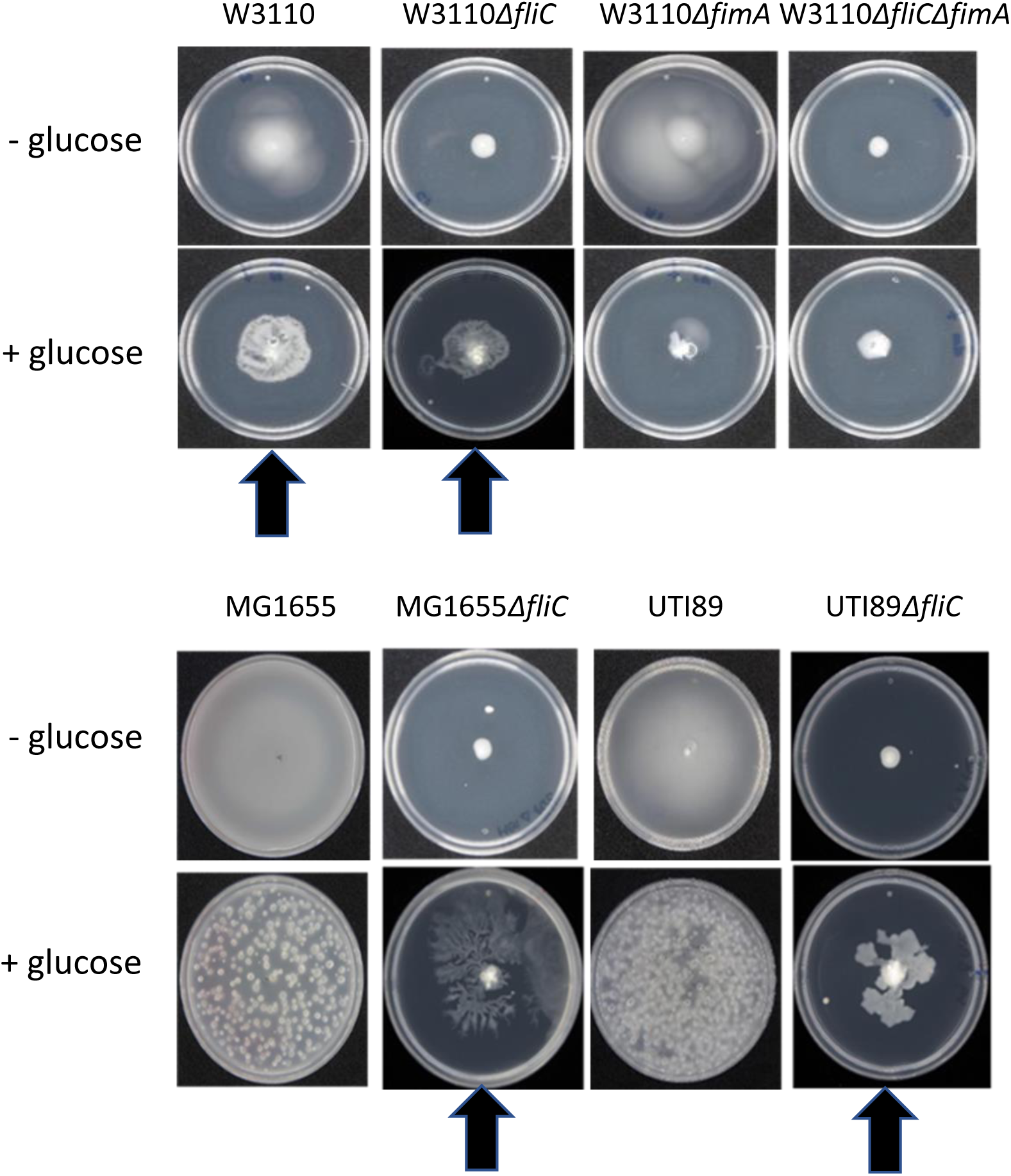
Motility observed in plates with 0.25% agar with and without glucose. Arrows indicate pili-mediated surface motility which is only observed in the presence of glucose. The bubbles for MG1655 and UTI89 with glucose resulted from CO2 generation of bacteria that swam into the agar.

MG1655 is a nonpathogenic strain of *E. coli* that, like J1, has an IS1 element 106 bases upstream from the *flhDC* transcriptional start site that increases flagella synthesis. MG1655 swam into 0.25% agar plates with or without glucose (Fig 8). Without glucose, MG1655 Δ*fliC* did not swim, but with glucose this mutant moved on the surface (Fig 8). J1 exhibited the same properties (not shown). The same phenotype for two different strains shows that properties are not strain dependent.

Pathogenic UTI89 swam into a plate with 0.25% agar, with or without glucose, and this movement required flagella (Fig 8). Swimming with glucose is unexpected, since glucose should prevent cyclic AMP synthesis which is required for flagella synthesis (30). However, we have shown that glucose does not prevent flagella synthesis in several pathogenic *E. coli* strains, which shows that the absence of a glucose effect is not strain dependent (11). UTI89 Δ*fliC*, which lacks flagella, moved a little on the surface in the presence of glucose (Fig 8). This surface movement required pili because UTI89 Δ*fliC* Δ*fimA* failed to move (11). In summary, glucose promotes surface movement in strains lacking flagella.

We examined surface motility with 0.45% agar for W3110, MG1655, and UTI89 derivatives with a deletion of *fliC* which forces these strains to move with pili. (Δ*fliC* Δ*fimA* double mutants of these strains do not move (11)). All three strains moved well with 0.5% glucose, but not with 0.5% glycerol (Fig 9A).

**Figure 9:**
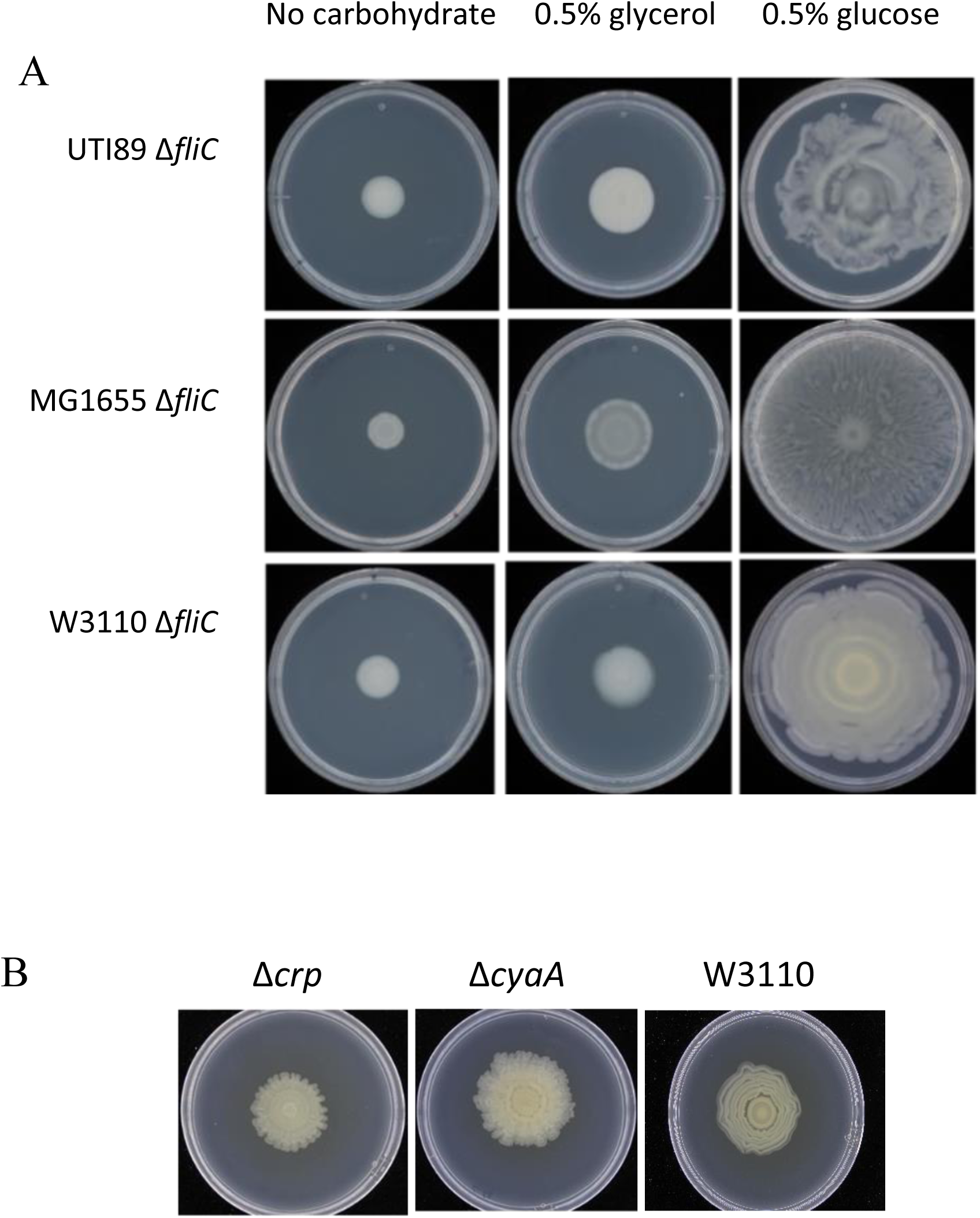
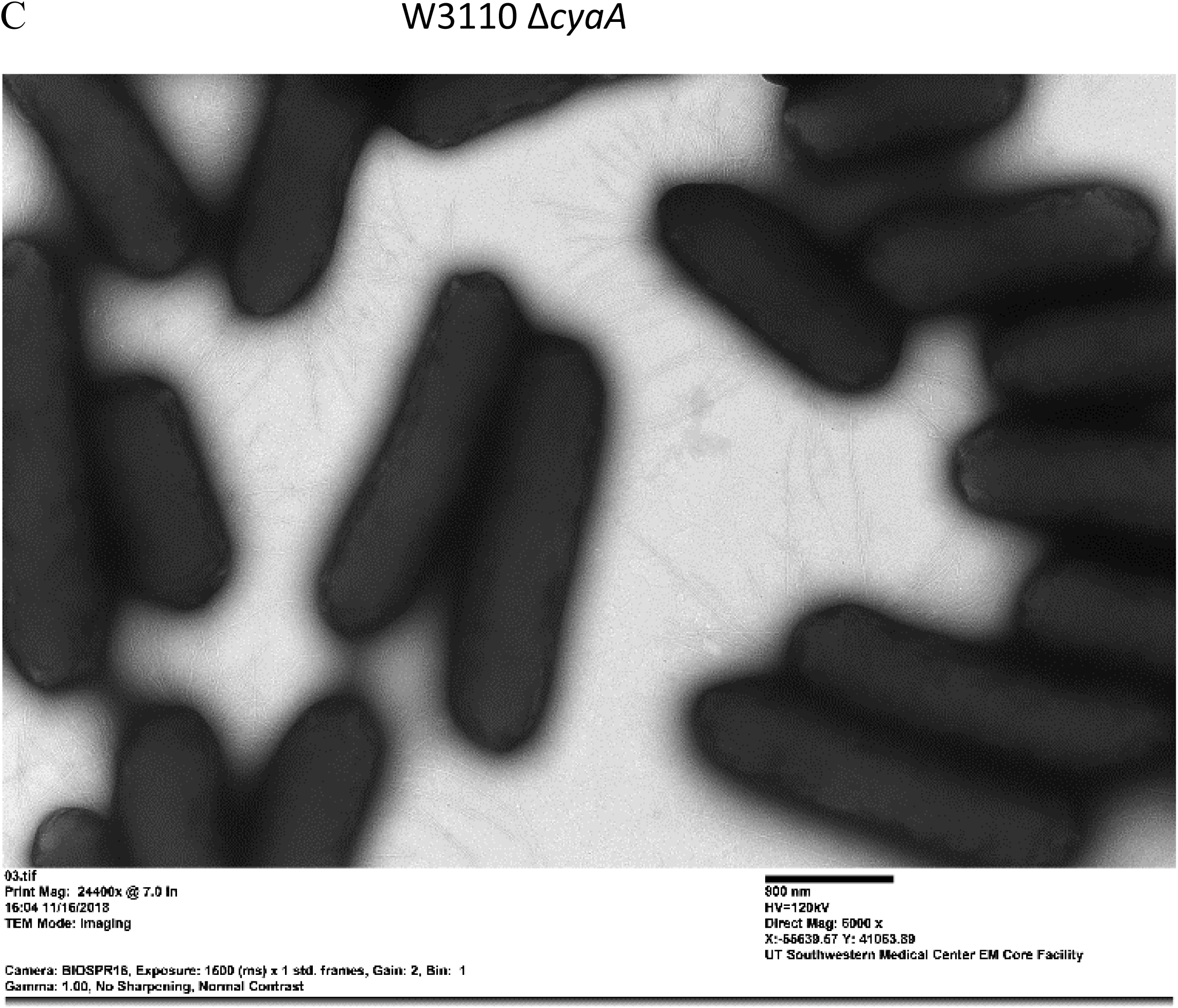
Pili-dependent surface motility of Δ*fliC* derivatives of UTI89, MG1655, and W3110. Δ*fliC* derivatives of UTI89 and MG1655 use pili in the absence of flagella, since Δ*fliC* Δ*fimA* derivatives do not move. W3110 normally uses pili for surface motility. (A) UTI89, MG1655, and W3110 mutants lacking flagella (Δ*fliC*) in medium with the indicated carbohydrate. (B) Surface motility of W3110 and Δ*crp* and Δ*cyaA* derivatives in surface motility medium with 0.5% glucose. (C) Electron microscopy image of the W3110 Δ*cyaA* mutant showing piliated cells. The pili are in focus, but the bacterial cells are not. The bar indicates 800 nm.

The glucose-dependent stimulation of movement predicts that pili-dependent surface motility will not require cyclic-AMP (cAMP), because glucose inhibits cAMP synthesis. Furthermore, CRP-cAMP represses *fimB* expression, which is part of the complex control of pili synthesis (31). Such control predicts that loss of CRP or cyclic AMP will not affect surface motility. As expected, *crp* and *cya* mutants of W3110 still exhibited surface motility, although the pattern of motility was altered (Fig 9B). Electron microscopy verified that the Δ*cya* mutant had pili, but not flagella (Fig 9C).

### The requirement for tryptone and the TCA cycle

Surface motility medium contains 1% tryptone. For all types of strains, 0.75% tryptone supported movement to the same extent as 1% tryptone (Fig 10). For J1 and UTI89, the bacteria on the 0.75% tryptone plate were not as dense as on the 1% tryptone plate, which suggests less growth (not shown). For all strains, 0.5% tryptone supported substantially less motility, and 0.25% tryptone did not support movement (Fig 10). The tryptone requirement does not distinguish between pili-dependent and flagella-dependent strains.

**Figure 10:**
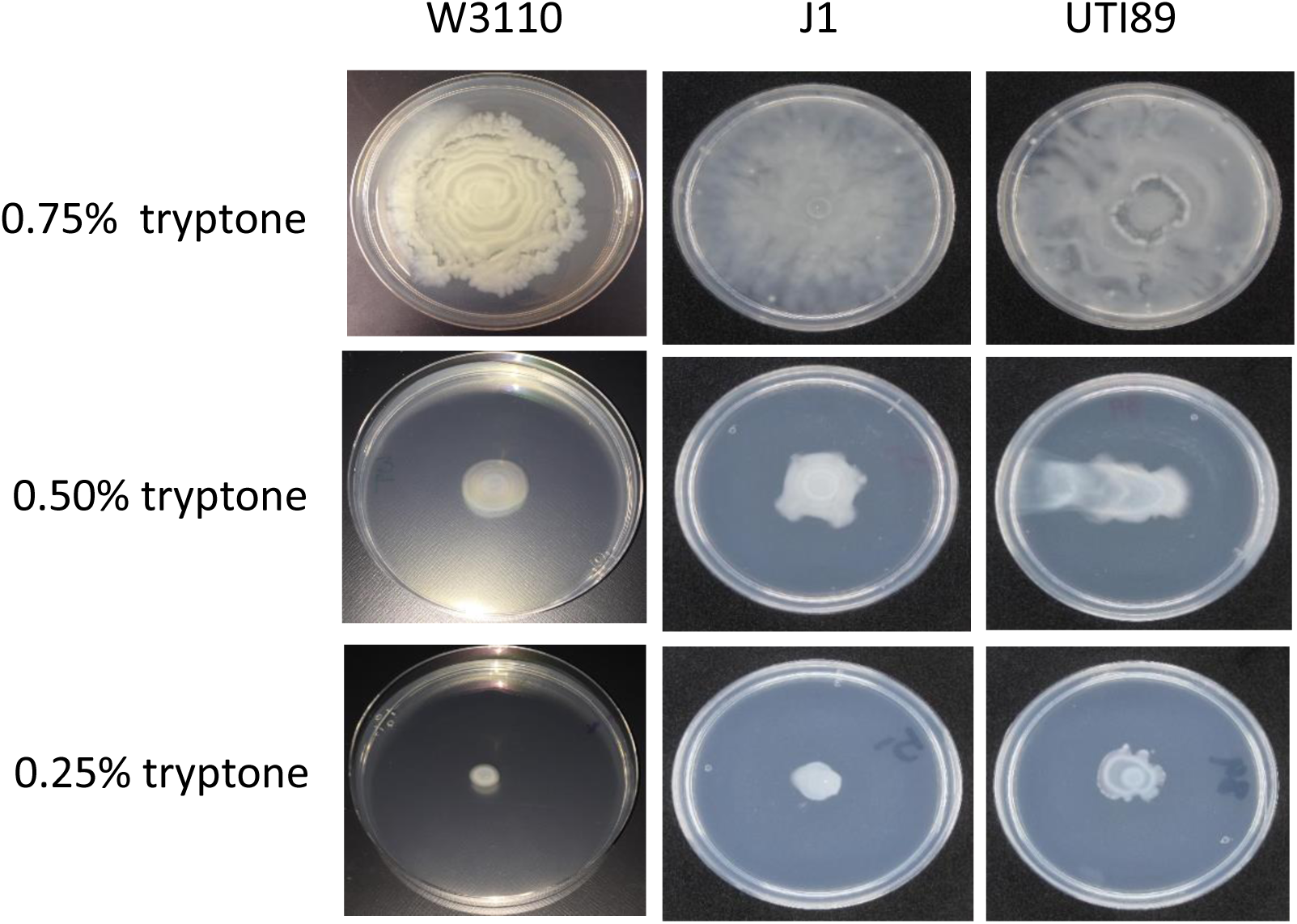
Tryptone requirement for surface motility.

Tryptone is an enzymatic digest of casein that consists mostly of amino acids, which can function as energy sources and biosynthetic precursors. Amino acid degradation in complex mixtures is poorly characterized, but if amino acids are energy sources, then their degradation requires the tricarboxylic acid (TCA) cycle, electron transport, and oxidative phosphorylation (32). Deletion of the following genes of W3110 had little or no effect on surface motility: *nuoC, glpD, glpA, poxB, hyaA, fdhF*, and *menA* (Fig 11). Movement was substantially impaired, but not eliminated, for mutants with deletions of *cyoA, ubiF, gltA, acnB, sucC, sucD, sdhA*, and *sdhB* (Fig 11). *lpd, sucA*, and *sucB* mutants could not move (Fig 11). The latter three mutants cannot generate succinyl-CoA, which is required for *meso*-diaminopimelate synthesis, an essential component of peptidoglycan that cannot be synthesized from components in the medium.

**Figure 11:**
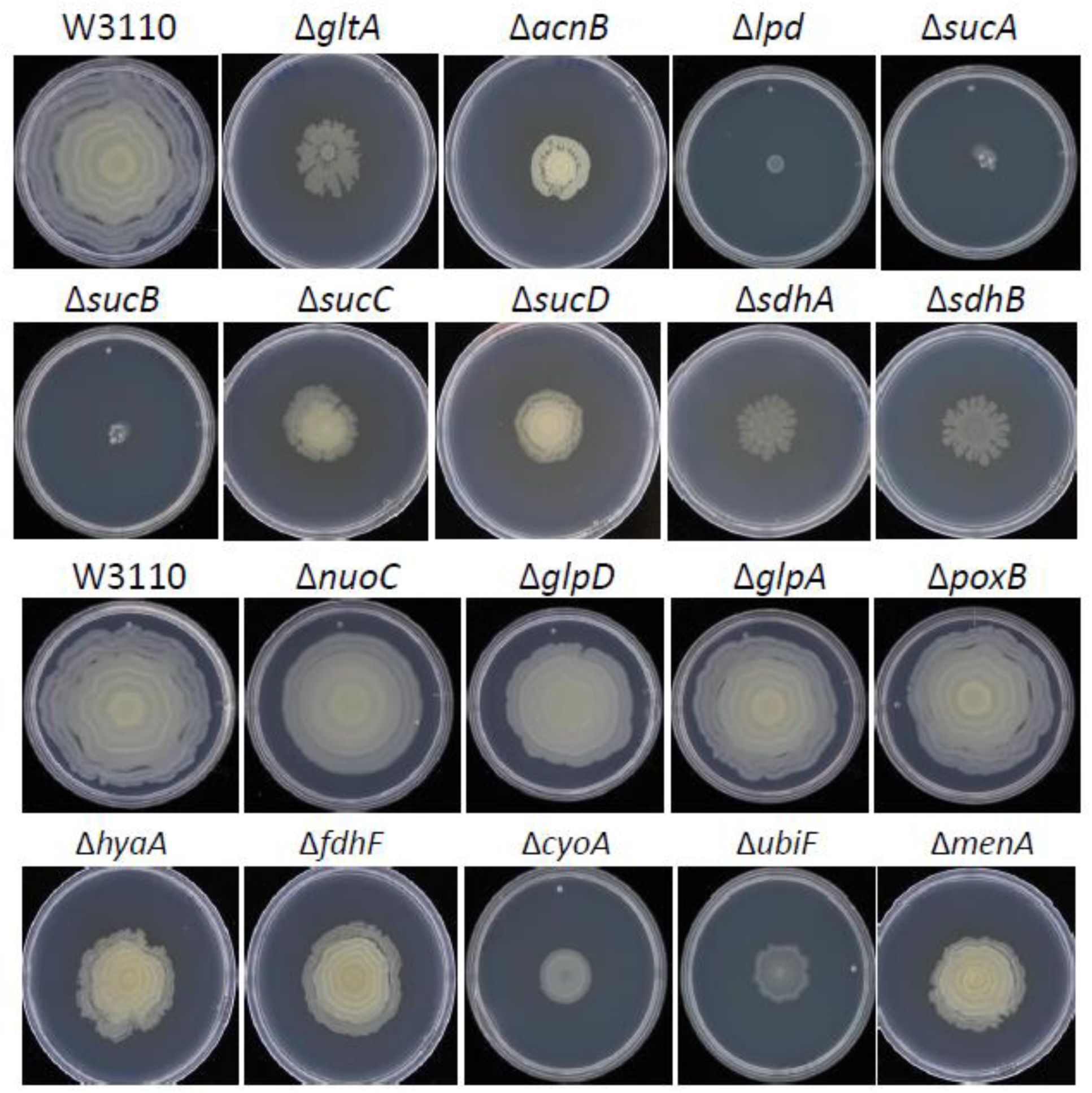
Surface motility of W3110 and derivatives lacking TCA cycle and electron transport chain enzymes.

For the flagella-dependent hypermotile J1 and uropathogenic UTI89, we examined mutants lacking genes for three representative enzymes of the TCA cycle: *gltA, sucC*, and *sdhA*. None of the three UTI89 mutants exhibited surface motility (Fig 12), which suggests that flagella-dependent movement requires the TCA cycle and oxidative phosphorylation. Such a result is not unexpected, since flagella rotation requires a proton motive force. In contrast, the J1 mutants moved, albeit poorly, and unexpectedly formed a ringed pattern like its parental W3110 (Fig 12). Such a pattern is more consistent with the pili-dependent movement of W3110, and electron microscopy showed that these mutants were piliated (Fig 12). These results suggest that a mutational block in the TCA cycle prevents flagella synthesis and results in pili synthesis in strain J1.

**Figure 12:**
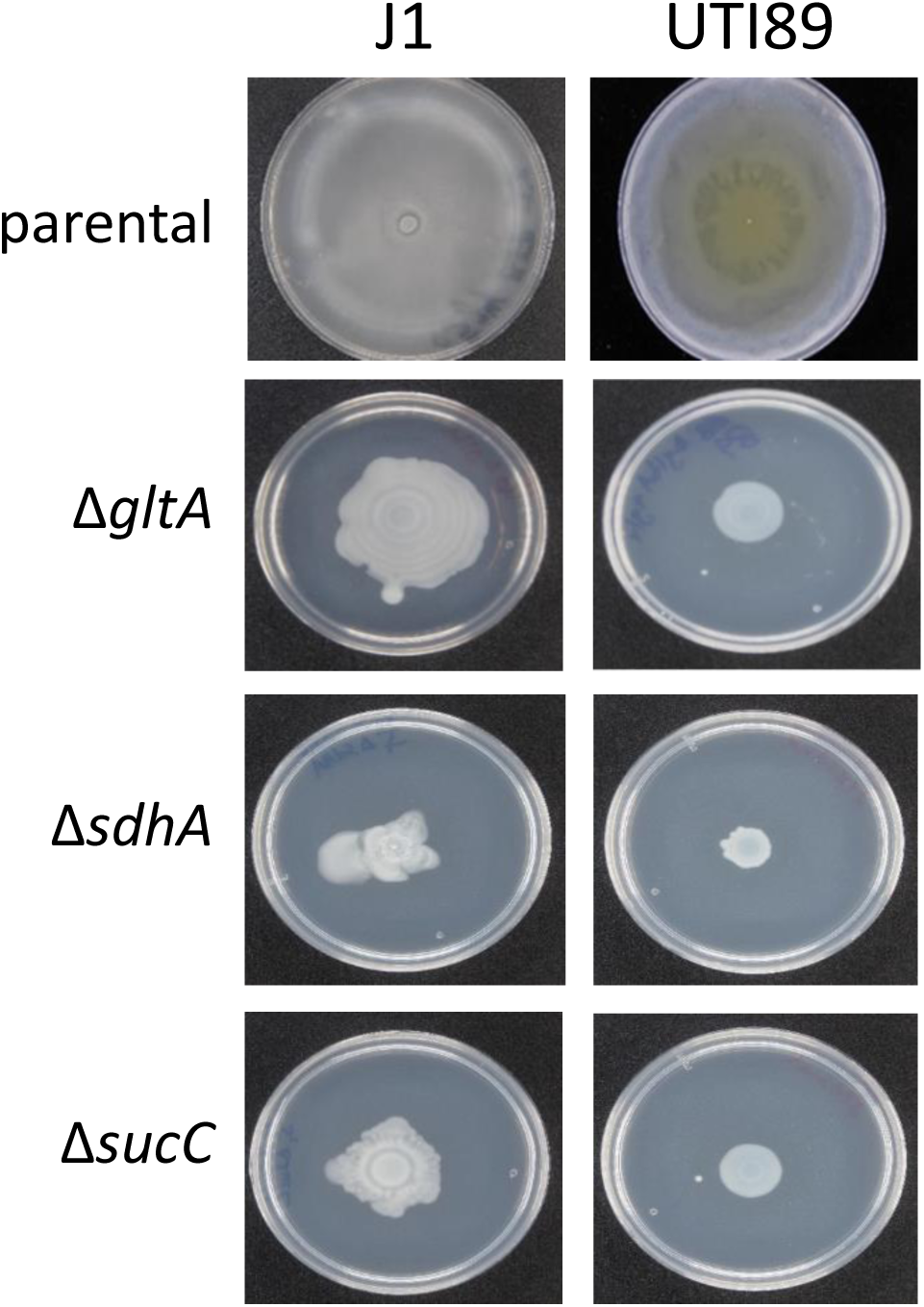
Surface motility of J1 and UTI89 mutants lacking representative TCA cycle enzymes.

**Figure 13:**
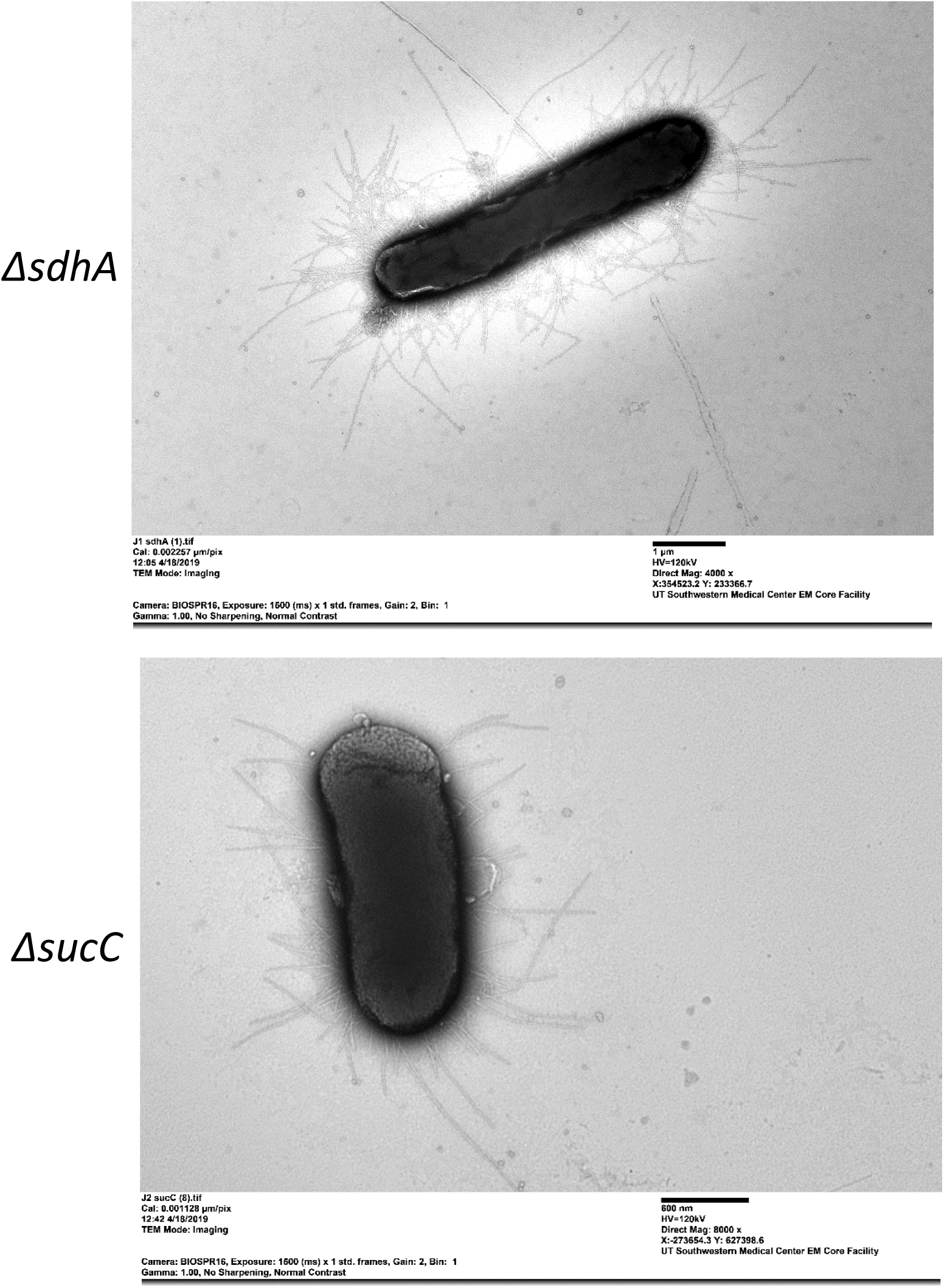
Electron microscopy images of J1 Δ*sdhA* and J1 Δ*sucC*. The bars for Δ*sdhA* and Δ*sucC* strains are 1 and 0.6 µm, respectively.

### pH of medium after motility assays

The pathway requirements for motility are clearly different between strains. Glycolysis through PfkA is required for motility of W3110 and J1, but not for UTI89. Loss of the TCA cycle affects all three strains, but the effect is greatest in UTI89. The motility medium is essentially unbuffered, which means that reliance on different energy-generating pathways will have different effects on medium pH: glycolysis will generate acids, and amino acid degradation via the TCA cycle will alkalinize the medium due to ammonia formation. The pH at the movement edge for W3110, J1, and UTI89 was 4.5-5.0, 5.5-6.0, and 6.0-6.5, respectively (not shown). The pH indicates the relative dependence of acid-generating carbohydrate degradation versus ammonia-generating amino acid degradation for these strains: W3110 is more dependent on glycolysis, whereas UTI89 is more dependent on the TCA cycle.

## DISCUSSION

Our goals were to determine the nutrient and pathway requirements for the surface motility of non-pathogenic *E. coli* that used pili or flagella for movement, and to compare these requirements with those of a uropathogen. The strains examined were W3110, which exhibited pili-dependent movement, and J1 and UTI89, which showed flagella-dependent movement. J1 is a hypermotile derivative of W3110. The results are summarized in Tables 1 and 2. *E. coli* pili-mediated motility has not been thoroughly characterized, but its requirements differ substantially from those for the flagella-dependent strains which confirms that pili-dependent *E. coli* surface motility is a distinct from flagella-dependent *E. coli* motility.

**Table 2:** Summary of pathway requirements for surface motility. a Mutants of J1 lacking TCA cycle enzymes moved partially and had switched from flagella- to pili-dependent movement.

| Function or pathway | W3110 | J1 | UTI89 |
| --- | --- | --- | --- |
| Glucose transport | PtsG | Not PtsG | Not PtsG |
| Embden-Meyerhof-Parnas (via PfkA) | Yes | Yes | No |
| Pentose-phosphate (via Zwf) | Yes | No | Yes |
| Entner-Doudoroff | Partial | No | Yes |
| TCA cycle | Partial | Yes/Partial <sup>a</sup> | Yes |

A potential problem with our analysis is the possibility that P1 transduction, which was used to construct the mutants, carried a mutation in a cotransduced gene that caused the phenotype. Complementation is used to address this issue, but control plasmids negatively affected motility. Instead, the conclusions are based on deletions of multiple genes in the pathways of glucose transport, glucose catabolism via the oxidative pentose pathway and Entner-Doudoroff pathways, acetogenesis, and the TCA cycle. In all cases, when loss of one enzyme of a pathway impaired movement, loss of other enzymes of the same pathway also impaired movement.

### Energy metabolism during surface motility

Flagella-dependent movement of UTI89 and J1 required the TCA cycle. For UTI89, the requirement for the TCA cycle was absolute, while for J1 the requirement appeared to be partial. However, the J1 mutants with TCA cycle defects used pili for movement. Our interpretation is that J1 flagella-dependent surface motility absolutely requires the TCA cycle, but under pressure to acquire nutrients can generate variants that utilize pili for movement.

Based on several observations, flagellar-dependent motility preferentially utilized amino acids degraded via the TCA cycle over carbohydrate degradation via glycolysis. First, movement of the flagella-dependent strains resulted in medium alkalinization, which can only result from deamination of amino acids. Second, both flagella-dependent strains had a lower glucose requirement than the pili-dependent W3110. Finally, J1 required fewer genes of carbohydrate transport and metabolism than parental W3110. Despite the greater reliance on amino acids and the TCA cycle, both J1 and UTI89 still required a carbohydrate.

Pili-dependent motility of W3110 was more dependent on carbohydrate degradation and less dependent on the TCA cycle, although mutants with defects in the TCA cycle were less motile. The evidence for this conclusion is (a) greater medium acidification for W3110 than for J1 and UTI89, and (b) defects in a greater number of glycolytic pathways affected W3110 motility. These results suggest that pili-dependent movement is more dependent on ATP from carbohydrate catabolism. Perhaps intracellular ATP can control the conformational states of type 1 pili, which could contribute to a form of motility (33).

In summary, flagella-dependent motility requires the TCA cycle, oxidative phosphorylation, and the proton motive force, while pili-dependent motility has a greater reliance on ATP from glycolysis. This conclusion is consistent with observations on *S. enterica* swarming cells which are morphologically and metabolically distinct with vegetative swimming cells (34). Although swarming cells require glucose, almost all enzymes of glycolysis were lower while several TCA cycle proteins were higher during swarming (34).

### Comparison of requirements for the two strains with flagella-dependent movement

The common requirements for J1 and UTI89 were glucose, albeit less glucose than the pili dependent W3110, and flagella synthesis in the presence of glucose, which is not a property of frequently studied *E. coli* lab strains. Despite these similarities, J1 and UTI89 also differed. J1, but not UTI89, required glycolysis through PfkA. In this respect, J1 is like parental W3110. Conversely, UTI89, but not J1, was affected by loss of the oxidative branch of the pentose cycle, the ED pathway, and acetogenic enzymes. Given their reduced requirement for glucose compared to W3110, carbohydrate metabolism may be important for one of more biosynthetic intermediate and the particular glycolytic pathway used for synthesis of the intermediate may not be important. For example, the specific pathway that generates triose-phosphates, e.g., the EMP vs ED pathway, may not be important if triose-phosphates are made. Another explanation for the differences between J1 and UTI89 is that the pathways used by the latter are an adaptation to the urinary tract milieu.

### Glucose transport during surface motility

Glucose transport in lab strains of *E. coli* requires PtsI (enzyme I), PtsH (the Hpr protein), and PtsG (the glucose-specific enzyme IIBC component). W3110 motility required all three components. J1 motility did not require any of these components, but still required a carbohydrate. A minor non-PTS glucose uptake system (e.g., GalP (35)) may be sufficient for J1’s reduced carbohydrate requirement. UTI89 did not require PtsG but at least partially required PtsH and PtsI. An additional glucose transport mechanism could explain the nonessentiality of PtsG in UTI89. PtsG-independent glucose transport could also account for flagella synthesis in the presence of glucose, if such a transport system does not control cyclic AMP synthesis. Glucose-independent flagella synthesis has also been observed for the UPEC strains PNK-004 and PNK-006 (11). PtsG-independent glucose transport and flagella synthesis in the presence of glucose, may be adaptations to the urinary tract environment.

### Comparing metabolic requirements for surface motility and UTIs

The pathway requirement for UTIs has been extensively studied with the uropathogen CFT073 in a competitive fitness mouse model (18, 36). The EMP pathway was dispensable for bladder infection but was required for kidney infection. On the other hand, CFT073 mutants lacking tricarboxylic acid (TCA) cycle, acetogenesis and gluconeogenesis enzymes were less fit in a murine model (18, 36). Like a CFT073 infection, UTI89 surface motility required the TCA cycle and acetogenesis, and did not require PfkA. However, a major difference is that a CFT073 infection requires gluconeogenesis (Pck), but UTI surface motility did not.

These and other results led to the conclusions that a CFT073 infection required amino acid catabolism via the TCA cycle, but not carbohydrate catabolism via PfkA (18). However, the requirements for UTI89 motility is consistent with a more complex explanation for CFT073 infectivity that accounts for some unusual phenotypes. While not requiring PfkA, UTI89 motility required a carbohydrate. This conclusion is based on failure to move without a carbohydrate and that only a derivative of UTI89 could move without a carbohydrate. Furthermore, some results from study of CFT073 infectivity also suggest that carbohydrate metabolism is important. While a *pfkA* deletion had no effect on competitive fitness in mice, a *pfkA pfkB* double mutant outcompeted the parental CFT073 strain in the bladder which suggests that glycolysis via phosphofructokinase is detrimental (37). Howeverm this result implies that carbohydrates are degraded. Furthermore, other evidence suggests that carbohydrate metabolism is important for UTIs (18). For example, genes for lactose and sorbitol catabolism are induced in intracellular bacterial communities during UTIs and their loss reduced UTI89 virulence (38), and loss of the Vpe carbohydrate permease, which transports an unknown carbohydrate, impaired virulence of the uropathogen AL511 (39). We propose that carbohydrate metabolism for both UTI89 and CFT073 occurs via PfkA-independent pathways. For example, loss of ED pathway enzymes results in defective UTI89 motility which implies that the ED pathway contributes to glucose metabolism. We note that during a UTI, bacteria will grow in urine, which contains low amounts of numerous carbohydrate that are not degraded via PfkA (18).

In summary, the metabolic requirements for UTI89 motility, which can be assessed in a defined and controlled environment, can provide insight into metabolism of uropathogens, including CFT073. The requirements for the TCA cycle and acetogenesis and lack of a requirement for PfkA for both UTI89 motility and CFT073 infectivity may suggest a UPEC-specific metabolism that is an adaptation to the urinary tract environment. However, UTI89 motility, but not CFT073 infectivity, requires the ED pathway, and CFT073 infectivity, but not UTI89 motility, required gluconeogenesis. These differences may be a function of either varied requirements for surface motility and infection or strain-specific differences. Strain-specific differences should not be considered surprising, since each UPEC strain must adapt to a different urinary tract environment. UPEC are becoming increasingly antibiotic resistant. The identification of UPEC-specific enzymes or processes, that do not show strain-to-strain variations, possibly a UPEC-specific glucose transporter, could be crucial for identification of targets for development of antibacterial therapies.

## MATERIAL AND METHODS

### Bacterial strains

*E. coli* W3110, hypermotile J1 (derived from W3110), NPEC strain MG1655, and uropathogenic UTI89 were used as parental strains. Table 3 lists the derivatives of these strains used in the study. The mutant alleles came from the Keio strain collection and contained deletions in which a kanamycin resistance gene replaced the gene of interest (40). The deletion/insertion was transferred by P1 transduction into the various parental strains (41). UTI89 was the uropathogen studied because of the ease of transduction. Other pathogenic strains can be transduced with P1, but the transduction procedure is more complex.

**Table 3:** List of bacterial strains

| Parental Strain | Mutants constructed | Parental source |
| --- | --- | --- |
| W3110 | $\Delta ackA$, $\Delta acnB$, $\Delta crp$, $\Delta cyaA$, $\Delta cyoA$,<br>$\Delta eda$, $\Delta edd$, $\Delta fdhF$, $\Delta fimA$, $\Delta fliC$, $\Delta fliC$<br>$\Delta fimA$, $\Delta glpA$, $\Delta glpD$, $\Delta gltA$, $\Delta hyaA$,<br>$\Delta lpd$, $\Delta menA$, $\Delta nuoC$, $\Delta pck$, $\Delta pfkA$,<br>$\Delta poxB$, $\Delta pta$, $\Delta ptsG$, $\Delta ptsH$, $\Delta ptsI$,<br>$\Delta sdhA$, $\Delta sdhB$, $\Delta serB$, $\Delta sucA$, $\Delta sucB$,<br>$\Delta sucC$, $\Delta sucD$, $\Delta ubiF$, $\Delta zwf$ | Lab Strain |
| J1 | $\Delta ackA$, $\Delta eda$, $\Delta edd$, $\Delta gltA$, $\Delta pck$, $\Delta pfkA$,<br>$\Delta pta$, $\Delta ptsG$, $\Delta ptsH$, $\Delta ptsI$, $\Delta serB$,<br>$\Delta sdhA$, $\Delta sucC$, $\Delta zwf$ | Lab strain (derived<br>from W3110) |
| UTI89 | $\Delta ackA$, $\Delta eda$, $\Delta edd$, $\Delta fliC$, $\Delta gltA$, $\Delta pck$,<br>$\Delta pfkA$, $\Delta pta$, $\Delta ptsG$, $\Delta ptsH$, $\Delta ptsI$, $\Delta serB$,<br>$\Delta sdhA$, $\Delta sucC$, $\Delta zwf$ | Lab strain |
| MG1655 | $\Delta fliC$ | Coli Genetic Stock<br>Center (Yale Univ) |

The kanamycin resistance gene was left in place, which implies the possibility of polarity. The conclusions of this work do not depend on whether gene expression is polar past the insertion. However, for several reasons, potential effects of polarity are likely to be minor. First, genes downstream from the insertion can be expressed from the resistance gene, although expression may not be the same as in the intact operon. Second, the insertion should not have downstream effects on the following monocistronic genes: *acnB, crp, cyaA, fdhF, fliC, glpD, gltA, menA, pck, pfkA, ptsG, ubiF*, and *zwf*. Third, insertions in *eda, lpd, pta, ptsI, sdhB*, and *sucD* are in the last gene of an operon and should not have a polar effect. Fourth, insertions in *cyoA, fimA, glpA, hyaA*, and *nuoC* were intended to eliminate a multiprotein complex: polarity is irrelevant. Finally, the insertion of the following genes may affect downstream genes but will only affect genes coding for proteins of the same pathway: *ackA* of the *ackA*-*pta* operon and acetogenesis, *edd* of the *edd*-*eda* operon and the ED pathway, *ptsH* of the *ptsHI* operon and carbohydrate transport, and the *suc* and *sdh* genes of their respective operons of the TCA cycle.

### Media and growth conditions

For growth on solid medium, strains were streaked on LB agar plates (10 g/l tryptone, 5 g/l yeast extract, 5 g/l NaCl, 15 g/l Difco agar) and incubated at 37°C for 15 h. For liquid cultures, bacteria were grown in LB broth with 25 µg /ml kanamycin (when appropriate) at 37° C with aeration (250 rpm) for 12 h.

### Motility Assays

#### Surface motility

Bacterial strains were streaked on an LB agar plate. After overnight growth, a single colony was inoculated in 1 ml of swarm medium: 1% tryptone, 0.25% NaCl, and 0.5% glucose and incubated at 37° C for 6 hr with aeration. Surface motility plates (swarm medium with 0.45% Eiken agar) were dried at room temperature for 4-5 hr after pouring. Changes to glucose and tryptone concentrations are indicated in the Results section. The motility plates did not contain antibiotics. One microliter from a 6 hr culture was inoculated at the center of the surface motility plate. Plates were placed in a humid incubator set at 33° C for nonpathogenic strains or at 37°C for UPEC strains, and surface motility was documented at 36 hours. Assays for the nonpathogenic strains were conducted at 33° C to ensure reproducibility: assays at 37° C were highly variable for NPEC strains because cells started moving at different times. Assays at 37° C for W3110 frequently result in generation of hypermotile variants. All assays were performed at least three times. Surface motility was extremely sensitive to conditions. Motility of the parental controls depended on the batch of the plates; for example, compare the results for parental W3110 in Figs 2 and 3. Motility of W3110 stopped if plates were removed from the incubator and examined for several minutes.

#### Swimming motility

Bacterial strains were streaked on LB, and a single colony was inoculated into 1 ml of swarm medium and grown for 6 h. Swim plates (1% tryptone, 0.25% NaCl, 0.25% Eiken agar) were stab inoculated at the center with 1 µl from the 6 hr culture and incubated at 33° C for 16 hr in a humid incubator. All assays were performed in triplicate.

### Electron microscopy

Cells from surface motility plates were collected from the edge of movement and fixed with 2.5% glutaraldehyde. Bacteria were absorbed onto Foamvar carbon-coated copper grids for 1 min. Grids were washed with distilled water and stained with 1% phosphotungstic acid for 30 s. 500-1000 cells were observed before choosing what to record. Samples were viewed on a JEOL 1200 EX transmission electron microscope at UT Southwestern Medical Center.

### PCR amplification of the *flhDC* promoter region

The *flhDC* promoter region was PCR amplified using FlhDp forward and reverse primers as described (42). The PCR product was then subjected to gel electrophoresis in a 0.8% agarose gel at 130 V for 30 minutes.

### pH of medium after motility

pH paper was placed directly on the plate.

## ACKNOWLEDGEMENTS

This work was supported in part by a UT Dallas Collaborative Biomedical Research Award grant program. The electron microscopy was performed at UT Southwestern which is supported by NIH grant 1S10OD021685-01A1.

